# An Open-Source Modular Framework for Automated Pipetting and Imaging Applications

**DOI:** 10.1101/2021.06.24.449732

**Authors:** Wei Ouyang, Richard Bowman, Haoran Wang, Kaspar E Bumke, Joel T Collins, Ola Spjuth, Jordi Carreras-Puigvert, Benedict Diederich

## Abstract

The number of samples in biological experiments are continuously increasing, but complex protocols and human experimentation in many cases lead to suboptimal data quality and hence difficulties in reproducing scientific findings. Laboratory automation can alleviate many of these problems by precisely reproducing machine-readable protocols. These instruments generally require high up-front investments and due to lack of open APIs they are notoriously difficult for scientists to customize and control outside of the vendor-supplied software.

Here, we demonstrate automated, high-throughput experiments for interdisciplinary research in life science that can be replicated on a modest budget, using open tools to ensure reproducibility by combining the tools Openflexure, Opentrons, ImJoy and UC2. Our automated sample preparation and imaging pipeline can easily be replicated and established in many laboratories as well as in educational contexts through easy-to-understand algorithms and easy-to-build microscopes. Additionally, the creation of feedback loops, with later pipetting or imaging steps depending on analysis of previously acquired images, enables the realization of smart microscopy experiments, featuring completely autonomously performed experiments. All documents and source-files are publicly available (https://beniroquai.github.io/Hi2) to prove the concept of smart lab automation using inexpensive, open tools. We believe this democratizes access to the power and repeatability of automated experiments.

## 1. Introduction

One of the core interests in modern life sciences is to understand how cells interact with each other and form highly organized biological systems. ^1,2^ Tools such as microscopy help to discover cell dynamics on very small scales in both time and space to provide evidence for previously formulated hypotheses. ^3^

It is common to repeat experiments many times or to use small variations in environmental parameters to understand causal relationships in increasingly complex biological systems. ^4^ This produces a lot of data, leading to challenges in data management and analysis. In most cases such experiments involve a lot of human labour carrying out protocols designed for and by humans, which is error-prone and might lead to data of lower quality and sometimes contribute to poorly reproducible results. ^5^

The resulting vastly improved datasets have the potential to lead to new insights. Here, a hypothesis is initially formulated and the experiment or parts of the experiment is designed and autonomously carried out by machines. ^2,6–8^ This is particularly useful when the same task has to be performed repeatedly over long periods of time, which is the case of multiple antibody-labelling. Supported by machine-learning algorithms, the machines (e.g. smart microscopes) are able to cope with the large amount of data generated in the process, as well as independently plan new experiments in order to arrive at even better findings, reduce the number of experiments and minimize the time required for simple tasks such as pipetting. ^9^ The application of machine learning algorithms in microscopy image analysis potentially helps to find correlations in data with previously trained networks ^10,11^ or in a completely unsupervised manner. ^12^ Notable are projects aimed at segmenting cells in micrographs tracking morphological changes, ^13^ improving the signal-to-noise ratio to enable low-photon dose imaging ^14,15 16,17^ or even performing augmented microscopy tasks.

The increasing complexity of experiments due to technologization, the associated cost of laboratory equipment, and the knowledge required to properly operate all the required instruments make high-throughput research very exclusive. ^18,19^ In turn, it is particularly challenging to train students to use such methods. ^20^

In the past, much open-source software and, more recently, open hardware projects have shown that by creating a community of developers and allowing them to develop the projects together, a high level of professionalism and usability is created, with the help being offered free of charge. In terms of the modern bio-lab, many research groups have already shown that different do-it-yourself (DIY) hardware can also be applied to biological questions and that it is even possible to build customized solutions using 3D printing and electronic setups, which significantly simplify the laboratory process. ^13,20–25^ Nevertheless, it is worth noting that the inhibition threshold for replication is still too high. After all, it requires a broad knowledge of electronics, 3D design and printing, programming and the subsequent frustration-dominated assembly of the devices. Even if open hardware is easily accessible, there is often a lack of high-quality documentation to finally replicate the result.

On the other hand, projects such as the 3D printed OpenFlexure Microscope ^23^, the Octopi/SQUID project ^13^ and the cellSTORM microscope ^22^ come with a fully illustrated manual that enables high-performance microscopy on a stand-alone or modular scale, as well as turning a smartphone into a Single Molecule Localization Microscopy (SMLM) setup. These projects have all been replicated by the growing community of microscopy users. For a great overview of the subject, please refer to. ^19^ In most cases, commercial devices lack open hardware or software interfaces for customization or automation, making it difficult or impossible to use them outside of the intended application for the devices, e.g., within a self-designed biological protocol. The Opentrons OT-2 pipetting robot ^6,26^ which we use in this manuscript is an example of commercially produced open-source hardware. Its commercial success shows that collaborative development also works in an industry context and additional functions from the developer community can be quickly incorporated.

Open-source software projects, such as the data analysis tool CellProfiler ^27^ or the image processing tool Fiji^28^, in which external developers draw on the core functionality, supplement it with their own Plugins and in turn make these available to others in online repositories with versatile application protocol interfaces (API), can accelerate the research process enormously by directly using the method or reusing code for their own research.

On close observation, such an interface between individual open hardware projects seems to be lacking at present. With UC2 we have already shown that it is possible to create a framework to connect different components from other projects or commercial sources so that a variety of optical experiments are possible. ^20^ Tools such as GitHub/Gitlab repositories in combination with proper open-source licenses help to organize such collaborations in the institutional context and beyond. This way, researchers and enthusiasts can participate and the lifetime of such a project very often outlives classical research projects.

In this work, we ask the question, how different expertise from several open-source hard- and software projects can be merged to generate a professional laboratory quality research tool. We show how the interaction between open-source tools such as the Opentrons OT-2 pipetting robot (Opentrons, New York, USA), the Openflexure Microscope (OFM Server ^29^), ImJoy and UC2, each expert in liquid handling, robotic microscopy, web-based image processing and modular optics solutions respectively, can help democratization of smart lab automation with a close-to turn-key solution. We will present two new compact slide-scanning (fluorescence) microscopes that can be placed directly in the Opentrons robot or cell culture incubators. Furthermore, we show an example of how a complete protocol from staining the microtubule network of fixed HeLa cells, to simultaneous in situ observation via the microscope, to quantitative analysis which can be realized via a simple interface in a web browser.

## 2. Methods

Here we give a brief description of the software pipeline, the automated pipetting system (**Figure 1a**) as well as the two different DIY high-throughput microscopy imaging systems (**Figure 1c, d**) for on-site microscopic imaging and image analysis inside the Opentrons pipetting robot (**Figure 1b**). Additionally, we show how all components can easily be integrated into a common workflow using Jupyter notebook to design complex biological protocols as depicted in **Figure 1a**).

**Figure 1.**
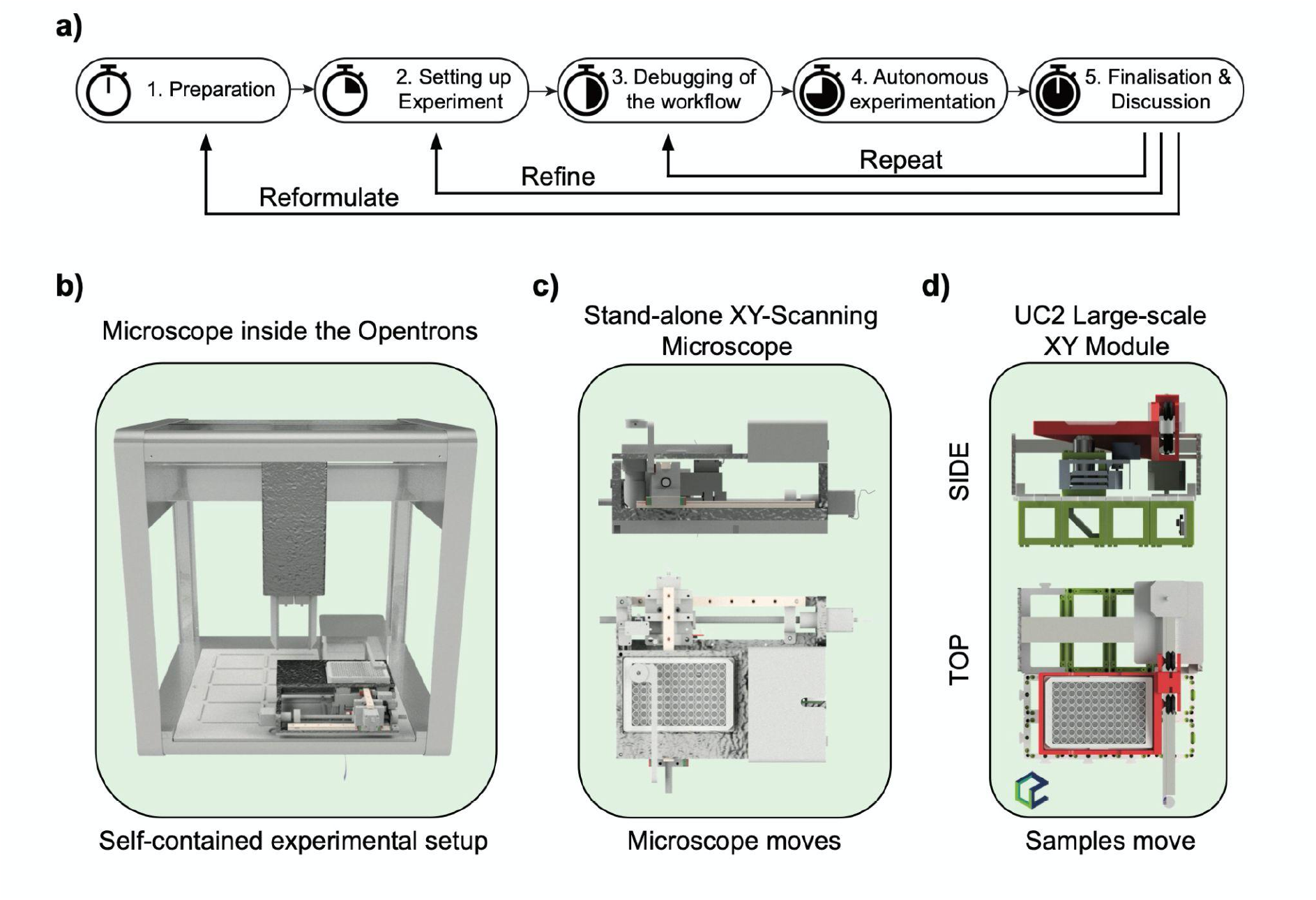
Components for a fully autonomous pipetting and imaging pipeline. a) A typical workflow starts with sketching the biological protocol and implementing it in python-based code for the opentrons/microscope. After simulating the code and setting up the experiment which typically involves placing reagents and samples inside the volume of the OT2, a first run can be started with some dummy reagents to debug the experiment. The autonomous experimentation includes robotic pipetting steps, continuous cell observation and processing on-the-fly image which can be controlled remotely. After collecting the data and an adjacent analysis, a hypothesis can be proven, the experiment repeated or refined. b) The microscope should fit into the working volume of the Opentrons OT2 pipetting robot to be integrated into a common pipetting workflow. b) The standalone high-throughput microscope “OpenMicronscope” moves the optical assembly around a fixed sample plate. d) represents an extension to the modular optical toolbox “UC2” and is based on a widely available laser engraving x/y table to realize high accuracy at a low price, where the sample is moved in x/y.

The first stand-alone microscope (**Figure 1c**) with a fixed sample stage was used as a development device for the software in several different locations (e.g. Sweden, Germany). From the experience gained from prototype development, the second UC2-based device (**Figure 1d**) moves the sample and was found much easier to reproduce.

### 2.1 Microscope Design

To perform cell observation both during the pipetting process and after completion of the protocol, we developed two compact microscopes with cellular resolution (~2-3μm) capable of recording time series of individual wells. The necessary parameters identified for the desired microscope for prototyping are:

1. flat design to fit into lateral flow hoods, biological incubators, liquid handling robots (i.e. Opentrons OT-2),
2. scanning of multi-well plates (6, 24, 96), as well as individual sample, slides with high speed and reproducibility with respect to sample coordinates,
3. transmission brightfield and optional fluorescence imaging,
4. easily reproducible, expandable, and cost-effective to quickly build a variety of instruments,
5. an easy-to-use control system that can be easily integrated into existing workflows,
6. simple integration of image processing tasks into the software,
7. perform in vitro experiments at temperature around 37 °C as well as high humidity for long periods of time with automatic focus.

Recent work such as the Incubot ^30^ or the open-source Lab Platform ^31^ has already shown how parts from a 3D printer or CNC machine can be used to create low-cost, high-throughput microscopes. We take up the ideas presented there and demonstrate two new compact 3D printed devices with a similar range of functions. They make optimal use of the limited working volume within the OT-2 and consider the maximum pipetting height of 150mm measured from the base.

The standalone “OpenmiTronScope” (**Figure** 1c, Supp. Material S1) offers a static sample well, moves the camera in XYZ to generate large-scale microscope images and uses a white-light LED to realize transmission brightfield imaging at cellular resolution. A more generic solution, where customized imaging schemes such as fluorescence or multiple objective lenses for different magnification and optical resolutions are desired, is provided by the UC2-based ^20,32^ system “Hi2” (**Figure** 1d). We extend the functionality of our recently introduced open-source modular optical toolbox UC2 with the capability of large-scale fluorescence microscopy with the development of a laser engraver-based XY stage (**Figure** 2a) and a novel Z-focusing unit (**Figure** 2b). Because most parts come pre-assembled, the complexity to replicate such devices is significantly reduced, hence enabling increased accessibility and wide-spread dissemination.

**Figure 2.**
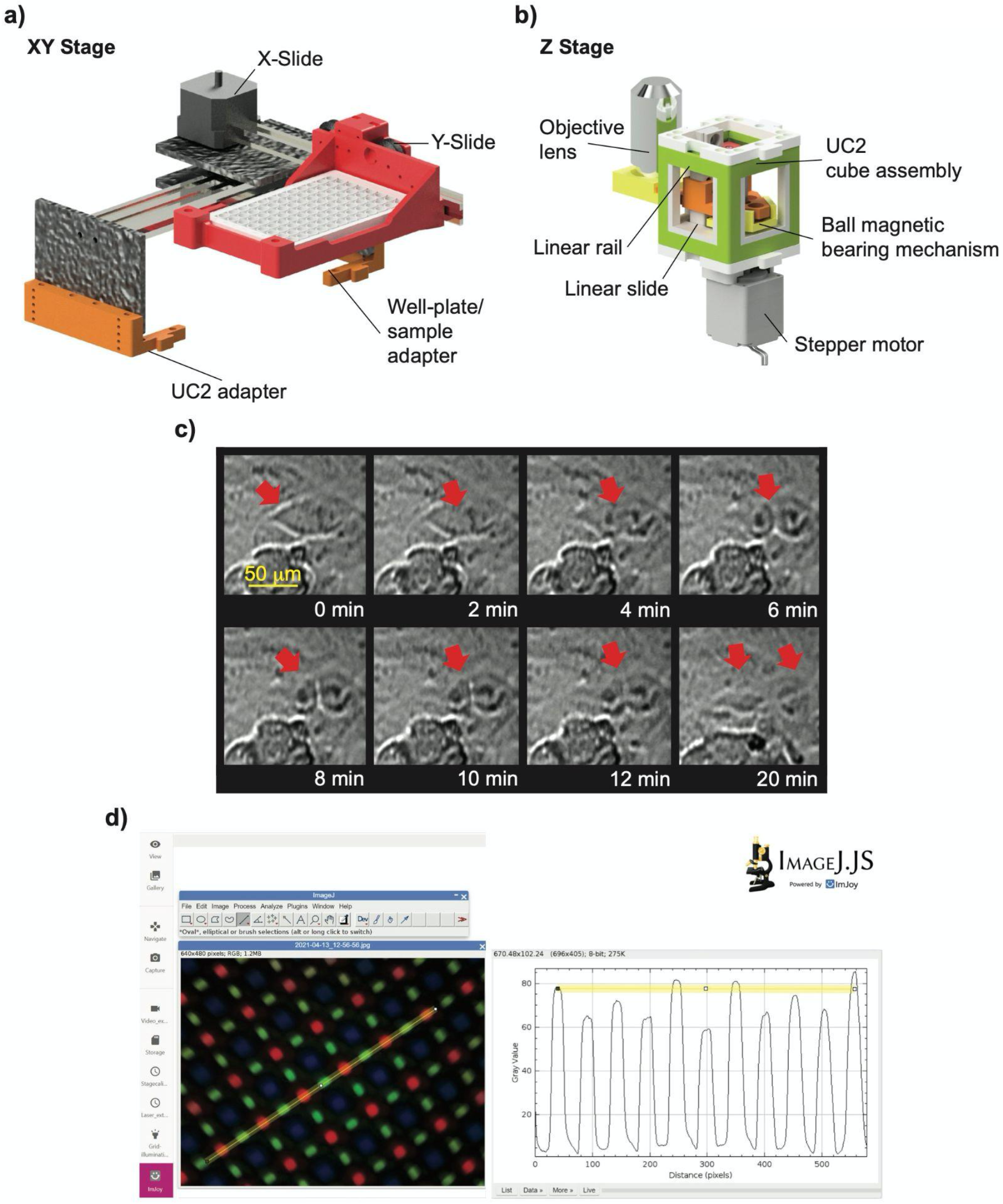
For the UC2-based high-throughput microscope “Hi2”, we developed two additional modules where the XY stage derived from a commercially available laser engraving machine can simply be integrated into the existing cube-based framework using a minimum of 3D printed parts. The novel Z-stage features low wobbling along the optical axis with the help of a magnetic ball-bearing decoupling mechanism in combination with linear rail bearings. The hardware module is optimized for simple replication, long lifetime and low price. c) A long-term time-lapse image series of in vivo HeLa cells conducted inside a cell culture incubator revealed a rare mitotic event. d) with the help of a newly created ImJoy extension for the Openflexure server, image processing, such as pixel size calibration using ImageJ.js, can directly be conducted inside the browser without transferring the data to external computers

An in-depth description and analysis of the mechanical properties can be found in the Supplementary Material (S1,S2), while a set of instructions on how to manufacture and assemble the parts as well as a detailed bill of materials (BOM) can be found on our project page https://beniroquai.github.io/Hi2.

### 2.2 Software for Hardware Control

For software control, it is important that all components, namely the pipetting robot, the image processing pipeline, the microscope and any additional components such as sensors or laboratory devices, can communicate with each other so that a fully automated workflow can be created. For this purpose, integration via a local area network (LAN) using Ethernet or WiFi, is a convenient option that allows the use of existing software to run each device, rather than requiring a single integrated application. Furthermore, intuitive control using a graphical user interface (GUI) or scripting interface such as Jupyter Notebook enables users to get started quickly with familiar tools.

In addition to an online protocol designer, the Raspberry Pi-based Opentrons OT2 can be controlled using a Python API from a Jupyter Notebook hosted on the OT2 or using a REST API via HTTP requests. ^26^ Similarly, the OFM server offers control through a browser-based GUI and using a REST API, which allows e.g. the initiation of scans and captures via the browser ^29^ or from a Python module running anywhere on the network.

The OFM server supports software extensions so that e.g. the “GRBL” interface ^33^ used here for the stepper motor control running on a serial-connected Arduino and the USB3 camera can be integrated easily. The server is designed for low computing resources and can be used on single-board computer (SBC) such as the Raspberry Pi (v3b, UK) or the Nvidia Jetson Nano (Santa Clara, USA). Because the GUI runs in the browser, the hardware can also be controlled remotely with suitable (i.e. secured) network configuration. Our core HTTP server code is available as the flask-labthings Python package. ^34^

### 2.3 Software for On-Site Image Processing

In order to process the generated image data directly without transferring each image from the microscope to an external processing machine or to involve imaging data into automated decision making for future pipetting events, we have integrated the browser- and web-based image processing tool ImJoy ^10^ directly into the OFM server. With this, it is for example possible to evaluate captured images directly in the browser using ImageJ.JS ^35^ or already available plugins such as the ITK/VTK Viewer. This is for example useful if the pixel size must be calibrated (as shown in **Figure 2d**), image tiles have to be stitched to form a larger field of view or time-lapse series have to be combined into a video. ImJoy also offers a large number of plugins developed by the community which can directly be accessed from within the browser. These include previously trained networks, denoising or deconvolution algorithms to partially compensate the loss of quality caused by the low-cost hardware. ImJoy can use the computing resources of the browser in which the plugin is operated, as well as external computer clusters, e.g. for training neural networks and is therefore not limited by the low computational resources by the SBCs.

## 3. Results

### 3.1 Open Collaboration for Distributed Hardware Development

The ability to develop hardware in a decentralized fashion has proven particularly helpful in the still ongoing COVID-19 pandemic, where many scientists have limited access to the lab. A discussion of an idea of building a compact microscope for the integration in the pipetting robot was followed by first prototyping of the “OpenMicrotron” in Jena (Germany). Two prototypes were sent to JCP and WO in Sweden at an early stage of development. With this functional prototype, it was possible to work on the further development of the software at several locations in parallel and to incorporate possible hardware and software optimizations into the next optimization iteration. Using tools such as remote coding sessions in VSCode (Microsoft, Seattle, USA), versioning tools such as Github/Gitlab and video conferencing such as Teams or ZOOM, the integration of the various components was thus realized from multiple locations even though the project partners never met in person. Any hardware changes can be quickly implemented using 3D printing or off-the-shelf components so that upgrades can be performed in situ. The result is the UC2-based system, where all issues from the first version have been resolved to maximize user experience and stability. This approach could serve as a blueprint for future projects since it offers a very fast development process from the first prototype in February to a fully working solution in May. Development was simplified by adopting an “openly developed hardware” ^18^ approach, removing the overhead of restricting access to designs.

### 3.2 Using the Hi2 Microscope for Live-cell and High-Throughput Imaging

For a high-throughput microscope, we identify the following criteria for use in biological experiments:

- Long-term imaging in the incubator at high temperature/humidity environment
- Permanent in-focus across the full well plate
- Reproducibility of the targeted XYZ coordinates over multiple runs overtime
- Ability to stitch a larger FOV from multiple image tiles
- Fluorescence imaging

3D printing material with increased glass transition temperature such as Polyethene terephthalate glycol (PETG) in combination with an autofocus algorithm, that compensates for temporal focus drift and tilted plates (see Supp. Material 3), helps to perform long-term imaging experiments of living organisms in cell culture incubators, where 37°C and high humidity represent challenging conditions for electronics and thermoplastics. A result for the use in incubators is shown in **Figure 2b**), where a rare mitotic event has been captured.

To estimate the mean displacement of multiple regions of interest within one well plate, a long-term experiment, where the microscope periodically (t=5 min) scans 32 out of the 96 wells, was performed at room temperature. The measurement in Supp. Material 3 suggests an average displacement of less than 32μm (4% for a region of interest) across an 18 h measurement, leading to the possibility to continuously monitor cells at multiple locations (e.g. wells). Fluorescent imaging (**Figure** 4) of the robot-labelled HeLa cell sample (Thermofisher alpha Tubulin Monoclonal Antibody cat. Number: A11126, Alexa Fluor 647^®^ Secondary Antibody, cat. Number: A-21235) the ability to stitch multiple image tiles to form a larger field of view directly on the device as shown in the zoomed region of interests (ROI) illustrated in **Figure 4**.

**Figure 4.**
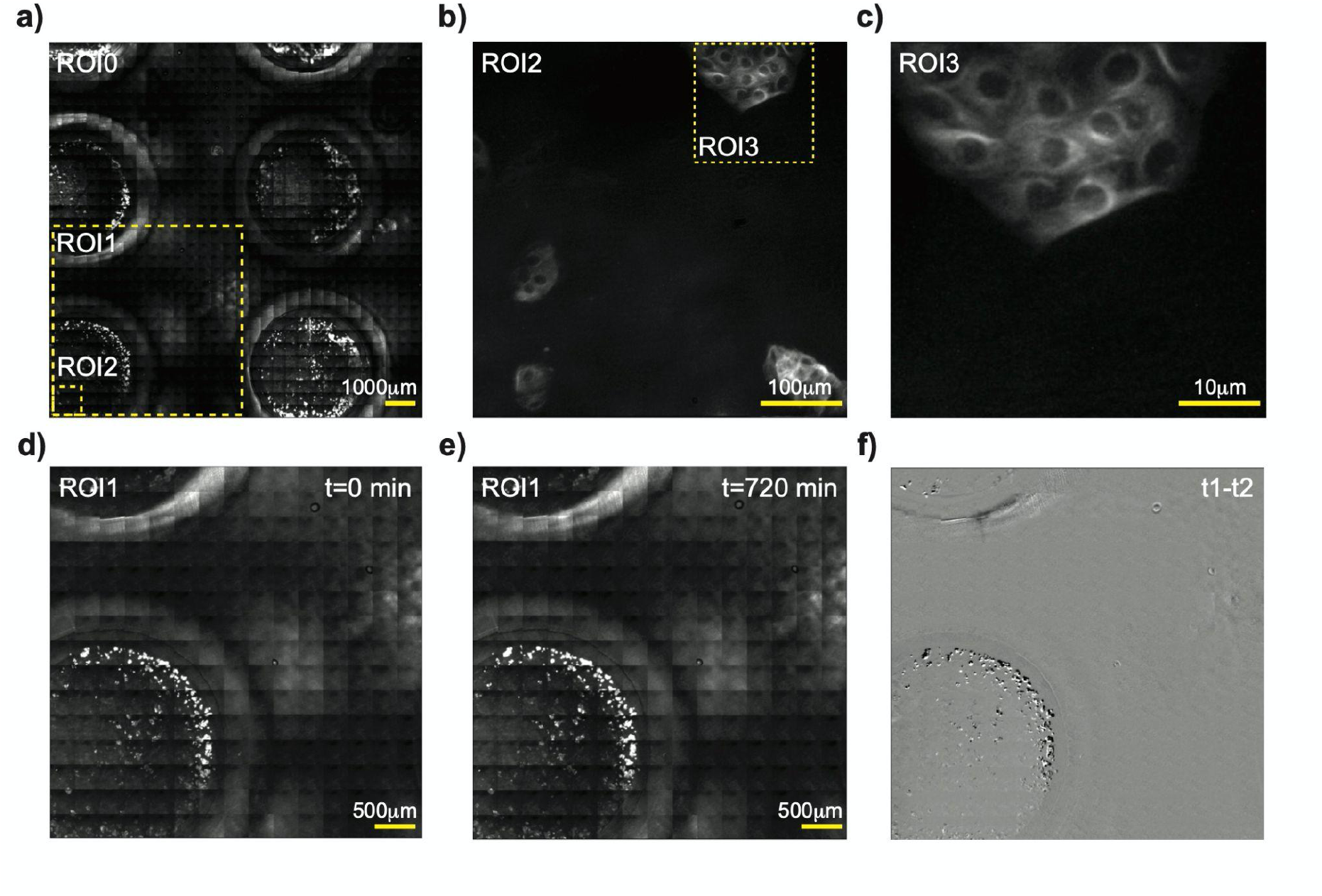
a) A representative stitched tile scan of a μSlide 15-well sample slide, where HeLa cells were labelled with anti tubulin AF647 primary/secondary antibodies using the Opentrons OT2. c) The zoomed-in Version (ROI3) of b) ROI2, suggests the presence of fibrous structures which can be identified as the microtubule network. d) - e) two roi-scans were performed 12h apart to demonstrate the location error computed in f as the difference between the two timestamps. Only small variations of the sample, mostly due to photobleaching can be observed.

### 3.3 Making the Labware Talk to Each Other

Until now, the individual components, such as the microscope, the GUI and the image processing worked independently from each other. In the following, we want to show how to connect them to create a fully automated workflow. This could in turn mean that pipetting steps can be done based on the previously obtained and processed imaging results to for example adjust pipetting volumes or have better environmental conditions for future experiments (e.g. buffer and antibody concentration).

A router (Netgear Nighthawk AC1900 R7000, 100€) ensures a stable connection of all devices over ethernet (Wifi for the Opentrons OT2) to ensure command as well as data transfer. An additional laptop or the Jetson Nano running the OFM Server are used to render the OFM GUI as well as the Jupyter notebook from the Opentrons. An in-detail description of how to set up the environment, perform simple tasks and a selection of ready-to-use pipetting protocols can be found on our project webpage (https://beniroquai.github.io/Hi2).

Using the Opentrons Python API, arbitrarily complex pipetting operations can be formulated in an easily readable code framework and executed both in standalone Python scripts as well as in dynamic browser-based Jupyter notebook. The latter has the advantage that each step is clearly logged, can be added to the lab notebook in a graphically easy-to-understand form, and facilitates reproducibility by sharing directly in the browser (**Figure 5**).

**Figure 5.**
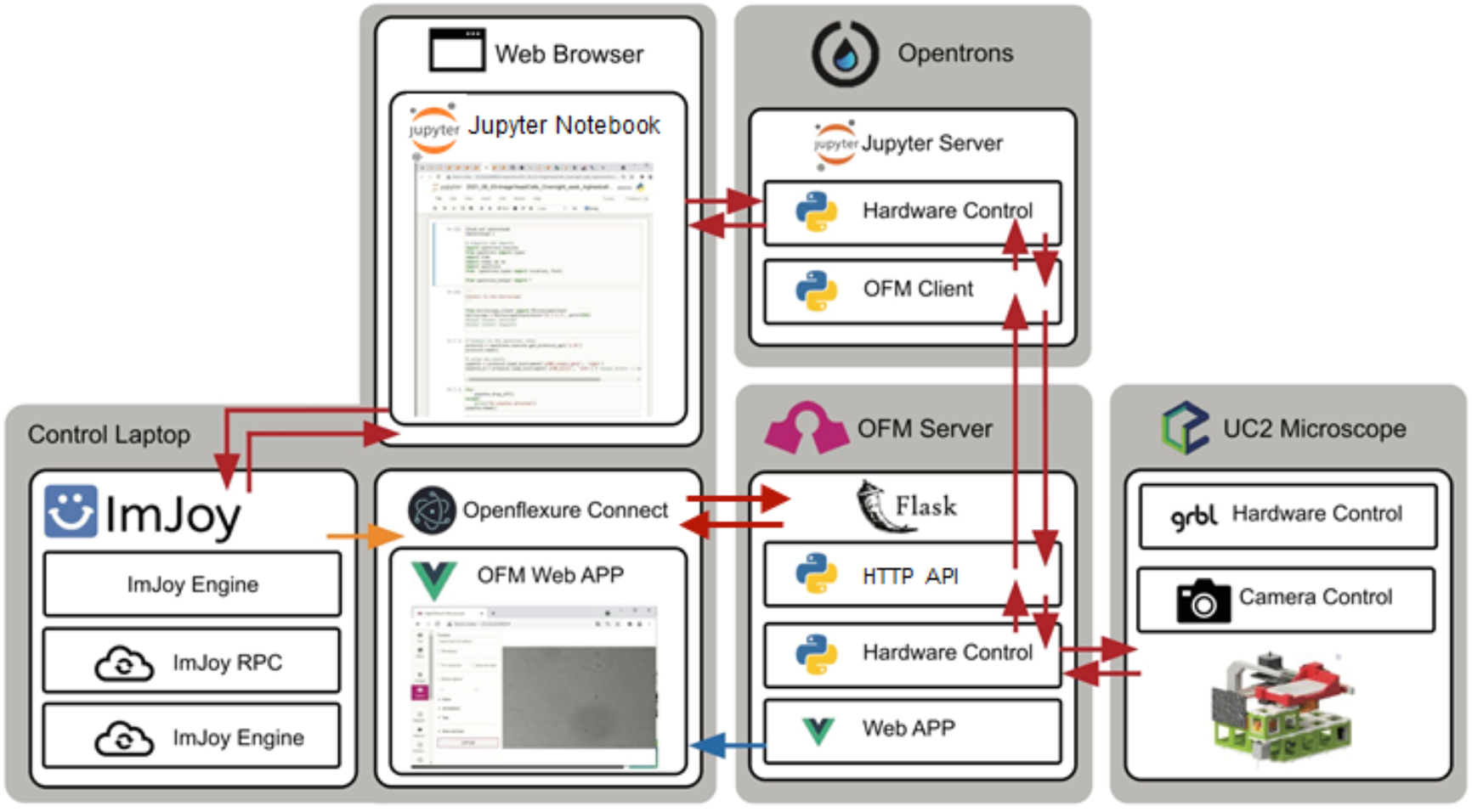
Diagram for intra-device communication, where the Opentrons OT2 pipetting robot’s Jupyter server controls the experiment. The protocol is formulated using human-readable python syntax, where on-the-fly image processing can be performed using available ImJoy plugins directly in the browser. Additional hardware, such as the GRBL stage and camera can be connected to the Jetson Nano, which also runs the OFM server.

The basic code structure for a protocol is divided into the definition of the labware (pipette tip rack, well plate, microscope, etc.), which are assigned to the deck coordinates and the execution of steps such as aspiration, dispensing or the movement from tip location A1 to well location B2 for example. To this end, common Python libraries (e.g. numpy ^36^) can be easily integrated, which is useful for controlling the Hi2 microscope or doing calculations based on computed results. Using the OFM client library (Figure 5, bottom) ^37^, Python commands such as image capture, move to coordinate or laser on/off are wrapped in human-readable protocol steps and executed via the REST API. This enables time-lapse imaging series during incubation times or fluorescence imaging after a staining process to check if the experiment worked on site. Custom labware, such as the microscopes or 3D printed tip racks, can be added using custom labware definitions or manually set XYZ coordinates.

The ImJoy Jupyter Notebook Plugin simplifies the execution of available plugins directly in the automated workflow. A typical use case is on-the-fly image processing, where the robot requests an image capture, downloads the data from the OFM server and transfers it to an externally running (e.g. browser of the laptop/computer cluster) ImJoy Plugin. This way, processing steps requiring high computational resources (e.g. segmentation by neural networks) can be realized without storing large data sets continuously. This enables the use of external libraries that cannot be installed on the read-only file system from the Opentrons and is especially useful for graphics rendering applications, where the ITK/VTK Viewer in the form of an ImJoy plugin helps illustrate results for further debugging.

Alternatively, for example in case an OT2 is not available, the microscope can conveniently be used as a standalone device, where ImJoy is integrated into the OFM web app, allowing for example image processing using ImageJ.js (**Figure 2d**) or other ImJoy plugins. The OFM GUI can be accessed through the browser or an Electron app, and summarizes the most important functionalities for common imaging tasks. An experimental feature offered by Opentrons enables remote control of the robot through its own HTTP interface. This enables the remote control of the robot from the OFM GUI, e.g. to move the LED mounted in one of the pipetting slots to the sample location. A brief description can be found in Supp. Material 4.

### 3.4 Imaging Results from a Fully Automated Antibody Labelling Workflow

The demonstration of a fully autonomously performed workflow is done by performing a classical, often very time-consuming laboratory experiment using primary and secondary antibodies for fluorescent immunostaining. As shown in **Figure** 6, the protocol consists of several pipetting and washing steps, with image acquisition during the waiting times to observe possible reactions of samples to reagents. The strength of such an autonomously working experiment is especially visible by repetitive runs with multiple different antibodies for different binding spots and subsequent imaging or bleaching of the stained molecules ^38^. These experiments can often last for several weeks. In the experiment presented here, only microtubules (anti-tubulin) are labelled with an Alexa Fluor ^®^ 647 conjugated (anti-mouse) antibody and then automatically imaged with the fluorescence unit (see **Figure** 6 right). After the image acquisition, the results are directly processed using ImJoy to apply a look-up-table (LUT), perform a pixel size calibration and quantify the cell’s features (see also Material S5).

**Figure 6.**
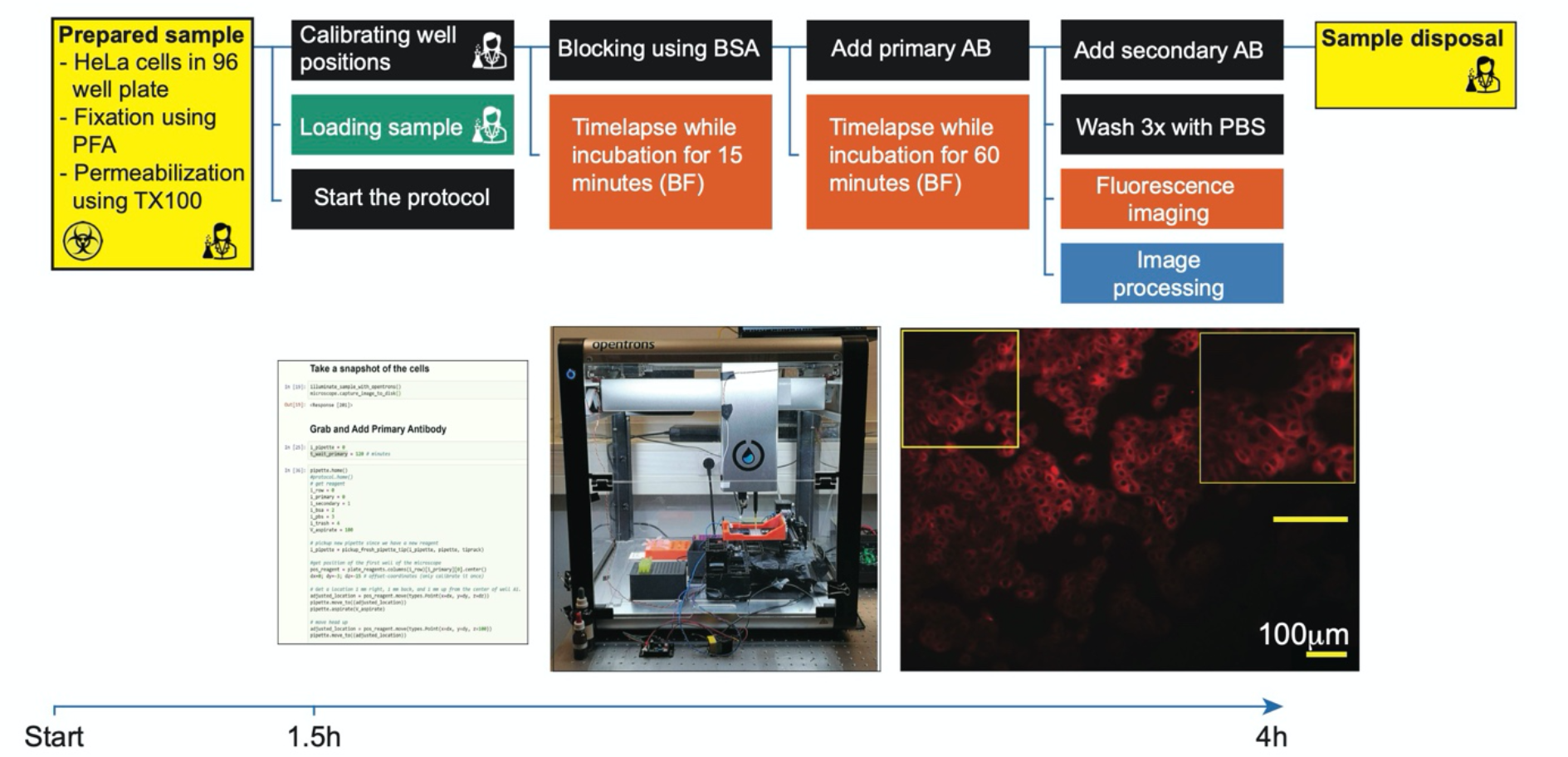
A common automated workflow that involves robot-assisted immunostaining, microscopic imaging and image processing. The control blocks are formulated in the form of a Jupyter Notebook running on the Opentrons OT2. All steps, except sample preparation involving fixation and permeabilization as well as sample disposal, are conducted directly at the robot.

A critical point of such an experiment is the debugging of the protocol without living samples and expensive reagents. For this purpose, the protocol can first be simulated before it is run on the robot without samples to detect and eliminate any errors in advance.

Live cell experiments inside the robot at 37°C ambient temperature and sufficiently high humidity are possible in principle. For this purpose, similar to ^39^, we have placed a hotplate heated to 75°C with a 2l water vessel in the working volume of the robot to achieve a constant temperature of about 35°C-37°C with the housing completely closed.

### 3.5 Performing “Smart Microscopy” by Finding Highest Cell-density

A unique feature of the closed-loop pipetting, imaging, and processing pipeline presented here is the ability to plan future steps within a protocol based on a computer-aided decision. We give a simple example in which the pipetting robot seeds an unknown quantity of yeast cells, which are imaged by the microscope after sedimentation (**Figure 7**). One image from each of the 96 wells is sent to a customized ImJoy plugin written in Python running on the laptop, which returns the number of yeast cells as the result. After cell preparation of all wells, the microscope moves to the position that has the lowest cell density and performs a long-time series. In this way, unknown cell densities could easily be calibrated, and subsequent experiments can be better selected in advance, e.g., to better observe the growth rate at the most ideal cell density without achieving over confluence, which can impact the biological outcome of a given experiment.

**Figure 7.**
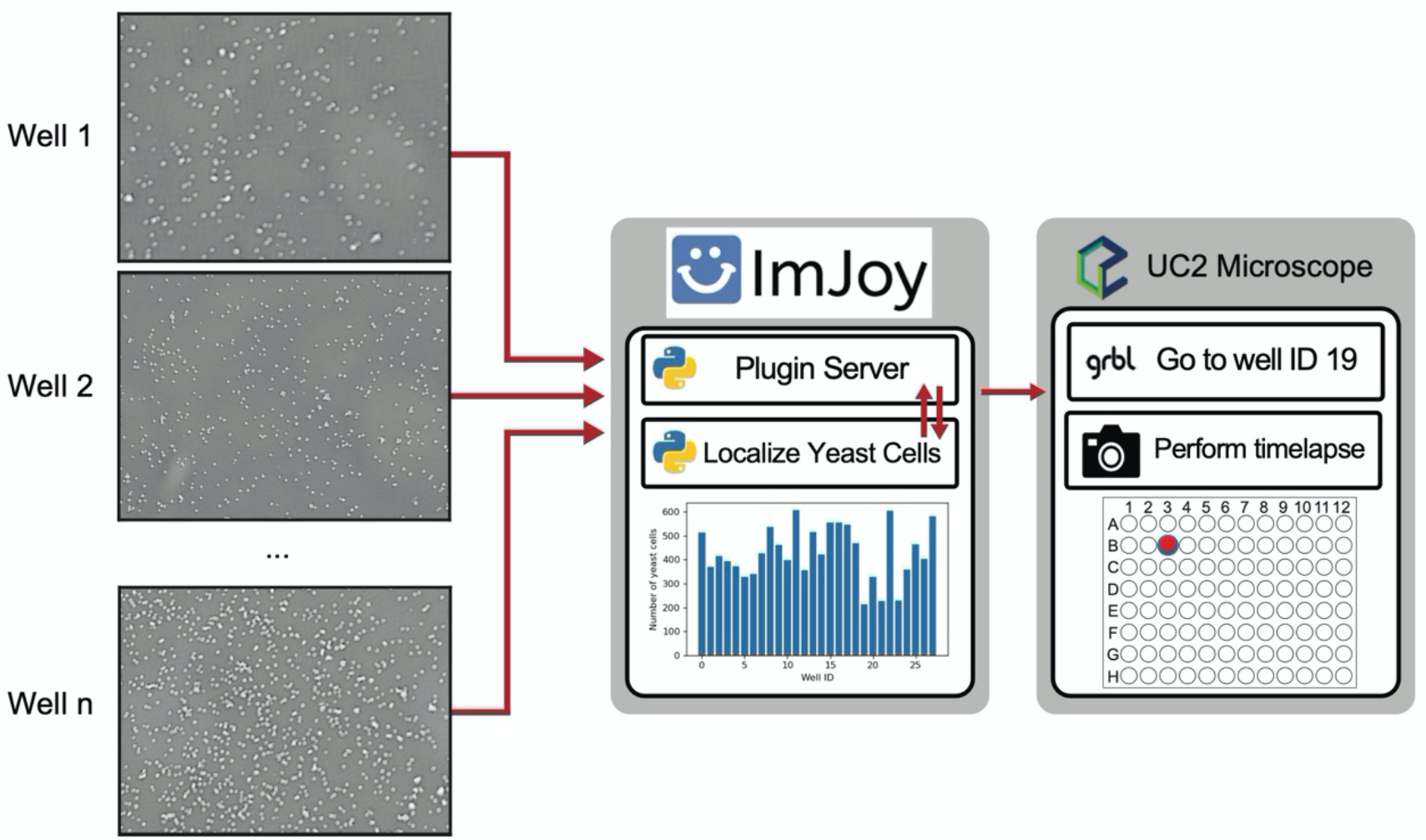
After seeding an unknown number of yeast cells using the pipetting robot, the microscope performs a whole plate scan before a dedicated ImJoy plugin running on a separated server computes the cell density and decides which well will be analyzed in long-time experiments. In this case well number 19 showed lowest cell density.

## 4. Discussion and Outlook

In this work, we demonstrate how a fusion of knowledge from open-source projects, each hosting a large and active developer community, leads to a powerful tool that makes the increasingly important field of lab automation available and accessible. The total cost, including the commercial OT-2, is about 7000€ and thus almost two orders of magnitude less than a comparable commercial setup with similar functionality. Additionally, it can be readily customized since the software and hardware can be adapted and extended to individual needs. The low price and the ability to work with it remotely also makes it ideal for high-safety lab environments.

Our solution includes browser-based GUI software for controlling the microscope, Jupyter notebook and ImJoy plugins for the reproducible execution of pipetting protocols and processing of image data. Imaging is performed on easily reproducible microscopes from the UC2 toolkit. This demonstrates that smart microscopy experiments can be realized without great effort and costs. We rely on existing projects and many off-the-shelf components like the commercially available open-source pipetting robot, laser engraving stages and pre-assembled UC2 modules to fully concentrate on the experiment.

The additional 3D printed parts can be easily adapted to the conditions of the experiment and replaced if necessary. However, this flexibility and the low price of the components are accompanied by significantly reduced stability compared to metal machined parts. In particular, the cantilevered sample holder in the Hi2 microscope is susceptible to low-frequency vibrations. This is particularly noticeable through small variations in the FOV and is especially visible when the robot is moving or when vibrations prevail in the building. 3D printed parts made from PLA start to soften at a temperature around 37°C (e.g. inside an incubator). Using PETG with a much higher glass transition temperature solves the problem, although a microscope warm-up phase with corresponding temperature-induced drift has to be considered. The low positioning error of less than +/- 4% within the FOV with repeated movements over 18 hours was a surprising finding for the low-cost x/y laser engraving table and shows the strength from the already well-engineered maker hardware for use in the scientific context. ^40^

Other critical points within an experiment are possible malfunctional behavior of components within an ongoing experiment leading to a cancelation of the process, as reagents such as antibodies are often very expensive and should not be wasted. This can be remedied by early debugging in the form of a protocol run with “dummy reagents” such as dyed water and checking the intermediate results or evaluating a series of time lapses made using a cell phone camera.

The low-cost optics used here are easy to obtain and can be replaced by e.g. higher quality lenses to achieve a higher optical resolution and better optical performance. Additional UC2 modules such as different excitation lasers and filter cubes can enable additional functionalities such as multicolour fluorescence imaging based on the building block-based principle.

The same applies to the here made integration of the web-based image processing tool ImJoy. Available libraries, code snippets and plugins for Fiji, Python or JavaScript can be integrated into the workflow, allowing the measured value in the form of 2D images of the image to be quantified and used for the next experiment or an adaptation of the previous workflow. However, a truly functioning feedback loop has not yet been established.

The open-source nature of all projects in this study pays off since they allow rapid integration and provide a legally valid framework through corresponding open-source licenses. The model of open, collaborative science we demonstrate here relies on the participation of many research groups. Often, direct commercial exploitation of research results still prevails, which can lock up intellectual property, and prevent projects like this one from using those results. ^41^

## 5. Conclusion

The effort to plan an already existing biological protocol, such as the antibody labelling shown here, and to transform it into a machine-readable code is currently much larger and more time-consuming than the manual approach. If the planning phase is included, the time required is an order of magnitude higher than conducting the pipetting manually. One can therefore ask the question of whom laboratory automation will be useful. However, with the workflows presented here making this technology available to a wide audience, the ability to share protocols that have already been performed, and the prospect of better training in this area already in the university context, it seems likely that automated performance of experiments will prevail in the long term. The ability to perform many experiments in parallel or replicate an experiment exactly will greatly improve both the quality of data and the reproducibility of studies. Our next goal is to perform closed loop experiments where, based on the data recorded in the experiments, trained neural networks can help to make decisions about future experiments and to plan and execute them autonomously.

## Acknowledgements

We thank Rainer Heintzmann for reviewing the draft of this manuscript. The authors want to thank Opentrons for supporting this project. We thank the Free State of Thuringia for funding BD. We acknowledge the ZIM project ZF4006820DF9 for funding HW. Additionally we thank the Royal Society (URF\R1\180153, RGF\EA\181034), EPSRC (EP/R013969/1, EP/R011443/1) for funding RB, KB and JC.

## Data Access Statement

All data raw and processed data associated with this manuscript will be accessible through the Zenodo **(DOI will be announced once the manuscript becomes officially published)** repository. All design files for the hardware, as well as manuals to install the software can be accessed through the GitHub repository https://github.com/openUC2/UC2-Hi2 or the project webpage https://beniroquai.github.io/Hi2.

