## Supplementary Information for "An Open-Source Modular Framework for Automated Pipetting and Imaging Applications"

Common laboratory tasks such as antibody labelling require trained personnel and typically many steps to conducting the protocol – an ideal job for a robot. However, telling a robot to simply move a liquid from a to b is already quite challenging. It involves the definition of multiple coordinates, commands to move the robot, processing the feedback from sensors to see if the task has been completed. In laboratory automation involving fragile live cell cultures, microscopic imaging, expensive reagents or incubation, the number of steps increases significantly. Nevertheless, the effort is worth it: The created protocol, in the form of a machine-readable algorithm/code (e.g., Jupyter Notebook) can easily be tuned and replicated, shared, and discussed, hence preserving and valuing the aspect of fully reproducible research briefly summarized in **Figure 1d**.
However, to have all components communicate with each other requires knowledge from many different disciplines, which is not always available. Our goal is to enable sophisticated laboratory automation in many laboratories by providing a simple recipe with open-source hardware and software components that helps to introduce common biological protocols in educational areas and bio/chemistry laboratories.

### S1 “OpenMitronscope”: Compact Standalone XYZt Well-plate Scanning Microscope

The optics module is equipped with a finite conjugate 10×, NA0.3 objective (Aliexpress/Newscope, 10€), where the folded optical path forms an image on a Raspberry Pi camera (v2.1, Sony IMX 219 sensor, 25€) with a final resolution of about 2.5 µm. Using a stepper motor driven flexure bearing, which reduces the linear motion of the driving screw by a factor of 3, the module can be precisely focused relative to the sample. A mechanism consisting of stepper motors (NEMA11, Eckstein, 18€), brass leadscrew (T12, China) and linear rails (MGN12H Slide/Rail, Roboterbausatz.de, 5€/15€) allows the lens and LED mounted above the sample to move in XY relative to the sample. For full operation, a Raspberry Pi 3b (UK, 50€), running the OFM server software as well as the Arduino-driven CNC shield v3 (China, 10€) were used. The total price is estimated to be between 300-400€. In-depth documentation can be found on the device’s project webpage (https://beniroquai.github.io/Hi2).

### S2 “Hi2” - High-Throughput Imaging Module for UC2

The recently introduced modular optical toolbox UC215 facilitates the prototyping of optical assemblies by reusing previously assembled modules such as lenses, mirrors, but also focusing devices and cameras. All optical, mechanical and electronic components are held in injection-moulded cubes (available upon request or alternatively 3D printed) with an edge length of 50 mm, which in turn are arranged on a base plate using a form-fit mechanism [32].
We take advantage of the toolbox's versatility and add a new focusing device with increased focus range and speed (**Figure** 2, right), as well as an XY scanning device that moves the well plate relative to the observing lens (Fig. 2 left). Compared to the previous setup, moving the sample has the advantage that more sophisticated optical setups can be integrated into such a system, which we will demonstrate with the addition of a fluorescence beam path implemented with a fiber-coupled laser diode.

For the XY stage (Fig. 2, left), we decided to incorporate a common design of a commercially available laser engraver (Neje Tools Laser Master Mini 2, China, 140€) with a scan area of 170×170 mm into the UC2 system. For this purpose, only two 3D printed adapters (10€) are needed to attach the stage to the injection-moulded cubes (13€ per unit). To mount the sample to the stage, the 450 nm laser supplied with the laser engraver is replaced by a multi-well plate holder that incorporates a spring-loaded lever mechanism to adjust the sample relative to the camera sensor plane.
The imaging path is an infinity-corrected microscope consisting of an objective (10× NA0.3, Aliexpress/Newscope, 30€) and a tube lens (Thorlabs ACM-100mm, 80€), with two kinematically mounted mirrors (Thorlabs silver protected mirror, 50€) to adjust the optical path. The newly designed focusing mechanism minimizes wobble, provides a fast linear movement, offers a low price and reduced complexity when assembling it. A worm drive realized using an M3 screw/nut translates an MGN12H linear ball bearing with a total focusing range of 25 mm and a theoretical step size of 1.25 µm to cover large sample tilts and correctly sample the depth of field also for higher NA objective lenses. In order to compensate for the global tilt we additionally perform a pre-scan of randomly selected points in the well plate and fit a 2D plane to them. The interpolated Z-coordinates for the focus give a very good estimate for the following fine-focusing. Important for a wobble-free operation is the decoupling of the drive axis from the motion axis. For this, we use a magnetic ball bearing mechanism, which decouples movement of the nut in XY due to imperfections in the screw from the motion of the objective. The setup consists of neodymium magnets located in the upper and lower half, respectively, supported by three 2 mm steel balls from one of the MGN12H bearings.

By adding a fluorescence filter cube (COMAR 740IY UK, 20€; Chroma 650/25, Germany, 250€) and a fibre-coupled laser (635nm, 150mW, 80€, Micost, China) which is first collimated by a plan-convex lens (Optikbaukasten.de, 5€) and imaged with a tube lens (165mm, CGI Shop, 10€) into the front focal plane of the objective, fluorescent imaging is possible (e.g. for Alexa Fluor ® 647). Adding additional laser sources and filter cubes realizes multi-channel fluorescence microscopy accordingly.
For the camera, we rely on a monochrome back-illuminated CMOS chip (Allied Vision, Alvium, Germany, 300€) which can be controlled and read out by the SBC Jetson Nano (Nvidia, CA, USA, 100€) running the OFM server. The microscope can also be extended with additional cameras or lenses to realize e.g. parallel colour imaging.
A white-light LED was mounted in one of the free pipette slots of the Opentrons to realize transmission brightfield illumination, where space restriction would otherwise make this technique hard to realize.
The Hi2-addon has a footprint of 200×200×180 mm, a total price of about 1000€, and can be fitted directly into the labware slots of the robot using screwable adapter plates.

Ouyang W. *et al.* **Fig. S1**

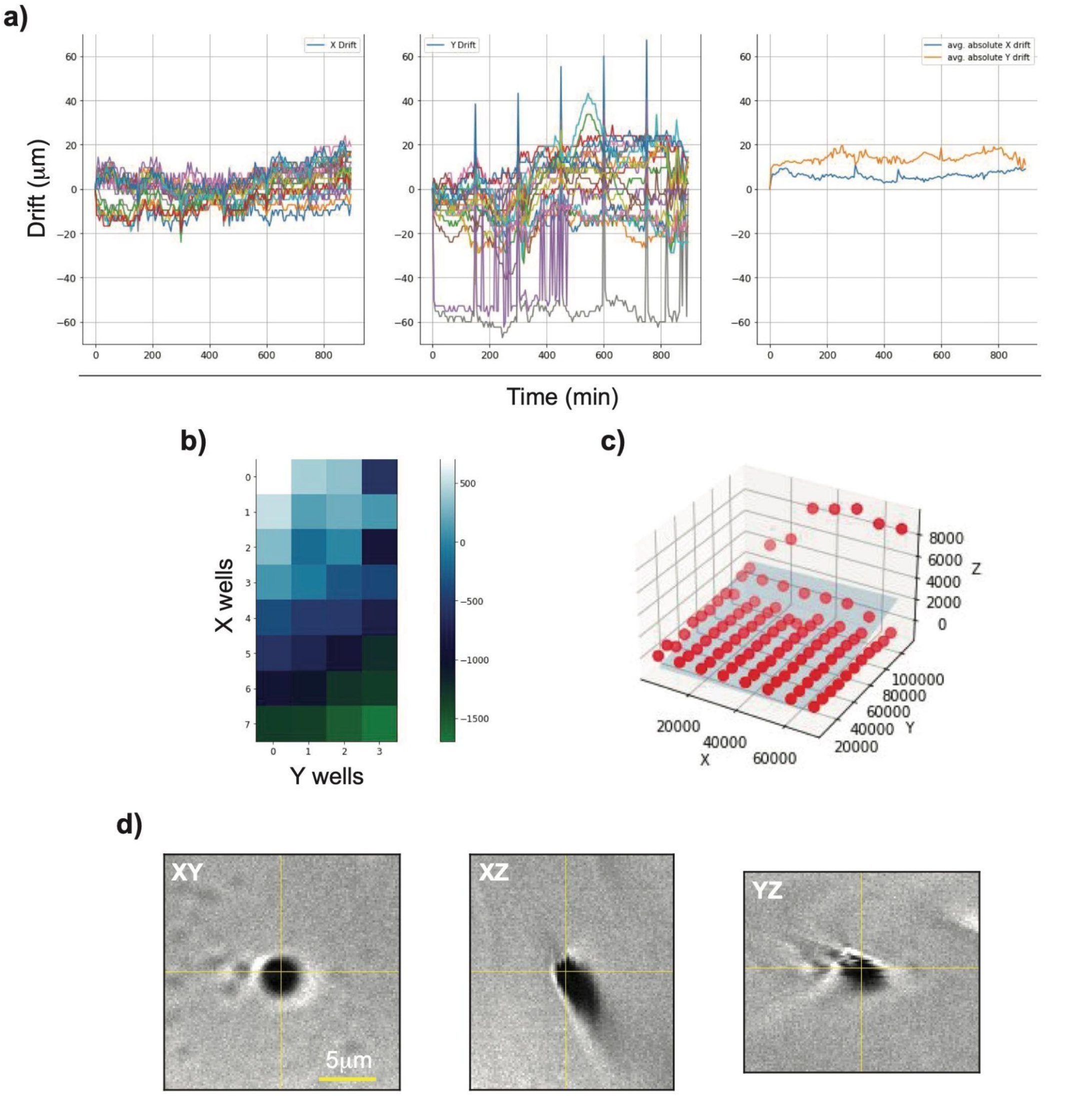


***Figure S1*** *a) the computed drift of repeated movements of 24/96 wells over 24 h shows a very small positioning error in x and y. b) the 96 well plate is mounted under an angle that can easily be precomputed by fitting a 2D plane to the points c) and compensated for with the integrated autofocus system. d) an exemplary 3D point-spread function shows only little wobbling along the Z-axis.*

### S3 Benchmarking the "Hi2" Microscope

We quantify the image quality and mechanical reproducibility of the UC2-based high-throughput microscope (Hi2) in a series of experiments.

For use in the incubator, where 37°C and high humidity represent challenging conditions for electronics and thermoplastics there were initially problems with the parts printed using polylactic acid (PLA), which already start to melt at these temperatures and contribute to significant focus drift. Polyethene terephthalate glycol (PETG) helps to reduce this effect significantly. Nevertheless, a residing temporal focus drift can be observed and was compensated for with autofocus (AF).

The extension-based OFM offers two different AF methods, where the first computes the image sharpness (e.g. Laplace/LoG filter) as a function of image slices along z and moves the objective to the maximum of a Gaussian fit of the sharpness plotted over z-position. The second and much faster method performs a full focal sweep while recording the size of the compressed JPEG stream. The slice with the highest file size (i.e. highest variation), usually corresponds to the best focus. Inconsistent timing of the focus stage's motion and the camera's frame acquisition prevented the latter from working consistently, hence we relied on the first method. An exemplary focus sweep of a scattering object in Fig. 3d) exhibits a slight wobbling along the y-axis visible by a periodic form of the 3D point spread function (PSF) along the optical axis. The x-axis, in contrast, only shows a constant tilt, which may be due to an uneven illumination relative to the objective lens.
The autofocus not only compensates for a temporal focus drift but also corrects for any tilt between the object and image plane respectively, which is shown in Fig. 3e), where the focus value is plotted as a function of x/y wells inside a 96 well plate. A long-term experiment at room temperature, where the microscope periodically (t=5 min) scans 32 out of the 96 wells and performs a refocusing every 2.5 h suggests very high reproducibility of the x/y coordinates over time. In **Figure 3b**), we plot individual x- and y-drift as well as the absolute mean position for all wells, which stays below 20 µm. while the maximum variation stays below +/-30 µm if outliers (e.g. due to lack of focus) are removed.
We continue this experiment by placing the device inside a benchtop incubator to observe in vivo HeLa cells in a similar experiment, where a scan of 24 wells was performed every minute with a refocus step every 15 minutes to compensate for the much larger thermal drift. This experiment allowed us to capture a rare mitotic event (Fig. 2b).

In addition to the transmission brightfield mode, the Hi2 setup also enables widefield fluorescence imaging using a fibre-coupled diode laser as presented in [13]. We test its performance using the automatically labelled microtubule structure of fixed HeLa cells. The static speckle pattern results in an inhomogeneous excitation pattern, which could be compensated with fiber shakers for example.
The high stage reproducibility enables simply concatenating the images without rearranging the XY coordinates, which enables the computation of a large FOV directly on the device as shown in the zoomed ROIs 1-3 in Fig. 4. We use this feature to again test the long-term stability by creating two large FOVs of the fluorescent samples (Ibidi, µSlide, Germany) 12h apart from each other. The computed difference shows only minor variation resulting mostly from bleaching of the sample.

### S4 Remote-controlling the OT2 from the OFM GUI

As an alternative to using the Opentrons as the central control, the OFM server can be configured as the central node of the workflow (**Figure 4**, top). Here, the control and image processing commands can be either formulated inside a Jupyter Notebook running on the Jetson Nano or with the help of customized OFM extensions.
To this end, we created a dedicated ImJoy extension accessible from the OFM GUI as exemplarily shown with the ImageJ.js integration run as an ImJoy plugin to calibrate the pixel size of the camera (**Figure 2d**). This helps to process captured images and image series directly in the browser and includes for example the fusion of multiple images to a time-lapse stack, a stitched larger FOV or multi-colour images.
Unfortunately, the ability to remotely control the Opentron via its existing REST API has been very limited so far. Although it is possible to access most - if not all - features remotely, the user-friendly Python API is not (yet) fully accessible to external clients like the OFM server. This includes access to the robot's state machine, which tracks the presence of a pipette tip or the various coordinates of lab equipment, for example. According to Opentrons, a more comprehensive integration of the HTTP API should be available soon.

In order to still have access to the most important functions, we have developed a customized client for the OT REST API that handles rudimentary commands such as moving the pipettes, and turning the light on/off, as well as loading and executing prepared protocols created with the Online Protocol Designer [26].
Using the ImJoy Jupyter notebook extension from within the Jetson Nano's notebook server, image processing tasks can be integrated into the workflow. The difference to the previously mentioned Scenario 1 is only the possibility to directly access the hardware of the Jetson by means of installed image processing libraries such as Tensorflow [42], OpenCV [43] or numpy [36] and thus no additional internet connection is required. Useful scenarios are e.g. the calculation of a camera overlay, wherein the live-preview e.g. the cell nuclei should be detected.
The integration of control commands into an OFM extension that acts as a dedicated GUI makes it possible to access useful functions, such as moving the brightfield LED mounted on the Opentrons directly above the sample, from the same browser window.

### S5 Comparing manual and robotic fluorescent labeling

Antibody-based labeling of cells is a very time-consuming laboratory task. The automated setup presented here can perform this task by reliably performing the same experiment repeatedly, but with different antibodies for different binding spots to increase throughput. Since the microscope is placed directly in the liquid handling robot, the sample is processed in situ, representing a clear advantage over a system in which the sample is placed at different locations to follow the steps of a given procedure. In Fig. S2a/b we compare the labeling results between a manually (**Figure S2a**) and automatically (**Figure S2b**) performed experiment, which resulted in a comparable microtubule staining. The Jupyter Notebook is similar to the manually performed antibody labeling protocol except for the fixation which was carried out outside the robot. It should be noted that the cells originate from two different batches. Both experiments, the manual and automated pipetting workflow, show equally good results where the filamentous structure can be vaguely seen on the micrographs taken with a 10x, NA=0.3 objective and a 635nm excitation laser. In automated pipetting, we found that pipetting depth is of great importance. A pipette tip that ends too high can result in an increased fluorescence background because the sample cannot be washed properly. A pipette that goes too deep into the sample may not be able to aspirate or dispense properly. Therefore, proper calibration in advance is required. The protocol is available in the Supplementary information.

Ouyang W. *et al.* **Fig. S2**

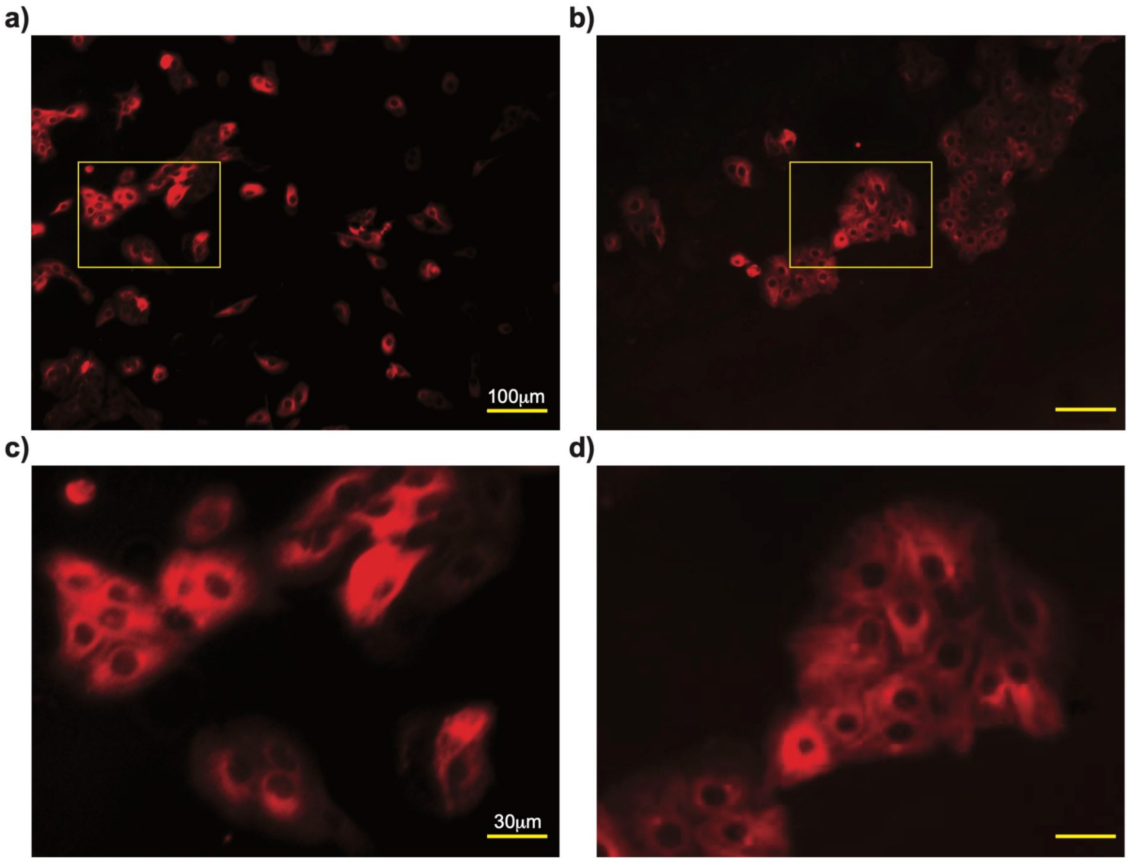


*Figure S2. We compare the automated a) and manual b) pipetting workflow for labeling microtubule filaments using a standard immunostaining protocol with primary anti-tubulin anti-mouse antibodies and anti-mouse Alexa Fluor ® 647 secondary antibodies. c) Zoomed imaging results acquired with the Opentrons-containing UC2 fluorescence microscope. The differences between automated and manual protocol might be explained be the need of further optimization of the automated protocol. d) In the case of the manual labeling protocol, the filamentous structure is clearly visible.*
